# Synthesis and Evaluation of Acridone and Xanthone Epoxides with Anti-MRSA and Anti-MSSA Activity

**DOI:** 10.1101/2021.12.10.472175

**Authors:** Padmaja Chittepu, Junshu Yang, Adam Benoit, Christine E. Salomon, Yinduo Ji, David M. Ferguson

## Abstract

A series of acridone and xanthone-based compounds bearing 1,2-epoxypropyl or 1,2-propanediol substituents were synthesized and evaluated for activity against MRSA and MSSA bacterial strains. The results indicate a correlation exists between the number of epoxide groups and activity, with peak MIC values observed for bis-epoxy derivatives. Both activity and heathy cell toxicity was shown to decrease with the addition of a third epoxy group. The corresponding ring-opened diol analogs were devoid of activity, demonstrating the critical function of the epoxide in mediating antimicrobial activity. The most active compounds were also screened using a regulated antisense RNA expression library. The results show no increase in activity against cells sensitized by down-regulation of the most common drug targets, including DNA gyrase, DNA topoisomerase, tRNA synthetase, and the fatty acid biosynthesis pathway. The compounds are postulated to function as membrane disrupting agents, similar to the xanthone natural product α-mangostin.

## Introduction

With the ever changing landscape of bacterial drug resistance, there is a pressing need for the development of new drugs with either improved activity against known targets or drugs with novel mechanisms of action that are less prone to the development of drug resistance. One of the most prevalent drug resistant bacteria that is of greatest concern is methicillin-resistant *Staphylococcus aureus* (MRSA). Known since the 1960’s, MRSA, which was once isolated to the health care environment, has now evolved and has become prevalent throughout the community.^1,2^ With our experience in the synthesis of acridone, xanthone, and acridine ring systems, we became interested in evaluating their potential as anti-bacterial agents.

Acridone alkaloids are a diverse class of natural products with the main source coming from the Rutaceae family of plants.^3,4^ While both naturally occurring and synthetic acridones have been tested and found to be active against cancers, herpes, psoriasis, and malaria, there has been little work regarding acridone based compounds as anti-bacterial agents.^5–8^ One recent study looked at a small sample of N-alkylated acridone products against both gram positive and negative bacteria which possess moderate anti-bacterial activity as compared to ciprofloxacin.^9^ Another recent study that coupled oxadiazoles to the acridone core produced compounds with moderate activity against a small sampling of gram positive and negative bacteria.^10^

In contrast to acridones, there has been substantially more anti-bacterial evaluation of naturally occurring xanthone alkaloids commonly found in Guttiferaeous plants, such as mangostins isolated from the mangosteen tree.^11–13^ Although mangostins have shown good activity against numerous gram positive bacteria as well as several drug resistant strains including MRSA, mangostins are also cytotoxic which has been attributed to their function as nonspecific cytoplasmic membrane disruptors.^14^ Despite the lack of a clear strategy to improve bacterial membrane selectivity, subsequent structural modifications to α-mangostin (shown in figure 1) to incorporate cationic groups that can interact with the anionic phospholipid head groups of bacterial membranes did result in substantially enhanced selectivity.^15^

**Figure 1.**
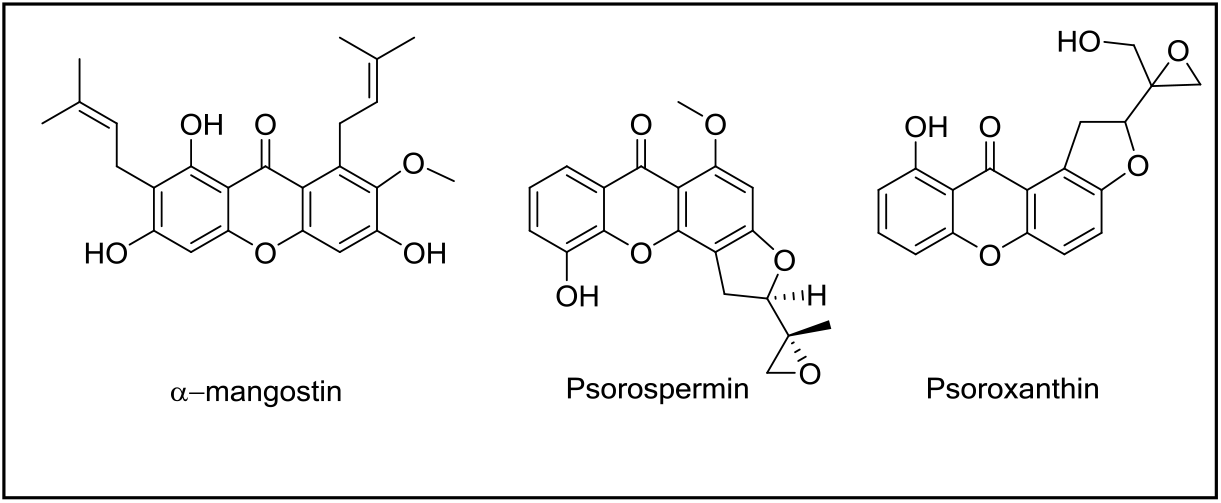
Xanthone natural products.

In this study, we explore a different approach to increasing the antimicrobial activity of xanthones and acridones by adding reactive epoxide groups to the ring system. Epoxy xanthones are found in nature as exemplified by the structures of psorospermin and psoroxanthin as shown in figure 1. In fact, the potency of psorspermin as an antileukemic agent has been attributed to the presence of the reactive epoxide that has been shown to alkylate DNA thereby inhibiting cancer cell proliferation.^16^ Herein we report the synthesis of several acridone and xanthone compounds bearing epoxypropyl substituents at varying ring positions. The ring opened 1,2-diol analogues were also synthesized as controls to probe the function of the epoxide. The compounds were evaluated for antibacterial activity using two MRSA strains (USA300 and USA400) and one methicillin-sensitve *Staphylococcus aureus* (MSSA) strain (MSA553).^17–19^ All compounds that were active against at least one strain of MRSA were also evaluated for Vero cell cytotoxicity as a measure of compound toxicity. Finally, a select sampling of active compounds were investigated using antisense RNA techniques to explore the potential mechanism of action.^20,21^

### Synthesis

Synthesis of the acridone-based compounds started with the condensation between phloroglucinol (**1**) and methyl anthranilate (**2**) to give 1,3-dihydroxy acridone (**3**), as shown in Scheme 1. Subsequent treatment of acridone **3** with cesium carbonate in a dimethylformamide/acetone mixture followed by epibromohydrin afforded the desired epoxy acridones **4-6**. By controlling the equivalents of epibromohydrin and base, we were able to synthesize all three analogs in two reactions. Hydrolysis of the acridone epoxides **4** and **5** to the respective diols **7** and **8** was accomplished in good yields by stirring overnight in 20% aqueous sodium hydroxide.

**Scheme 1:**
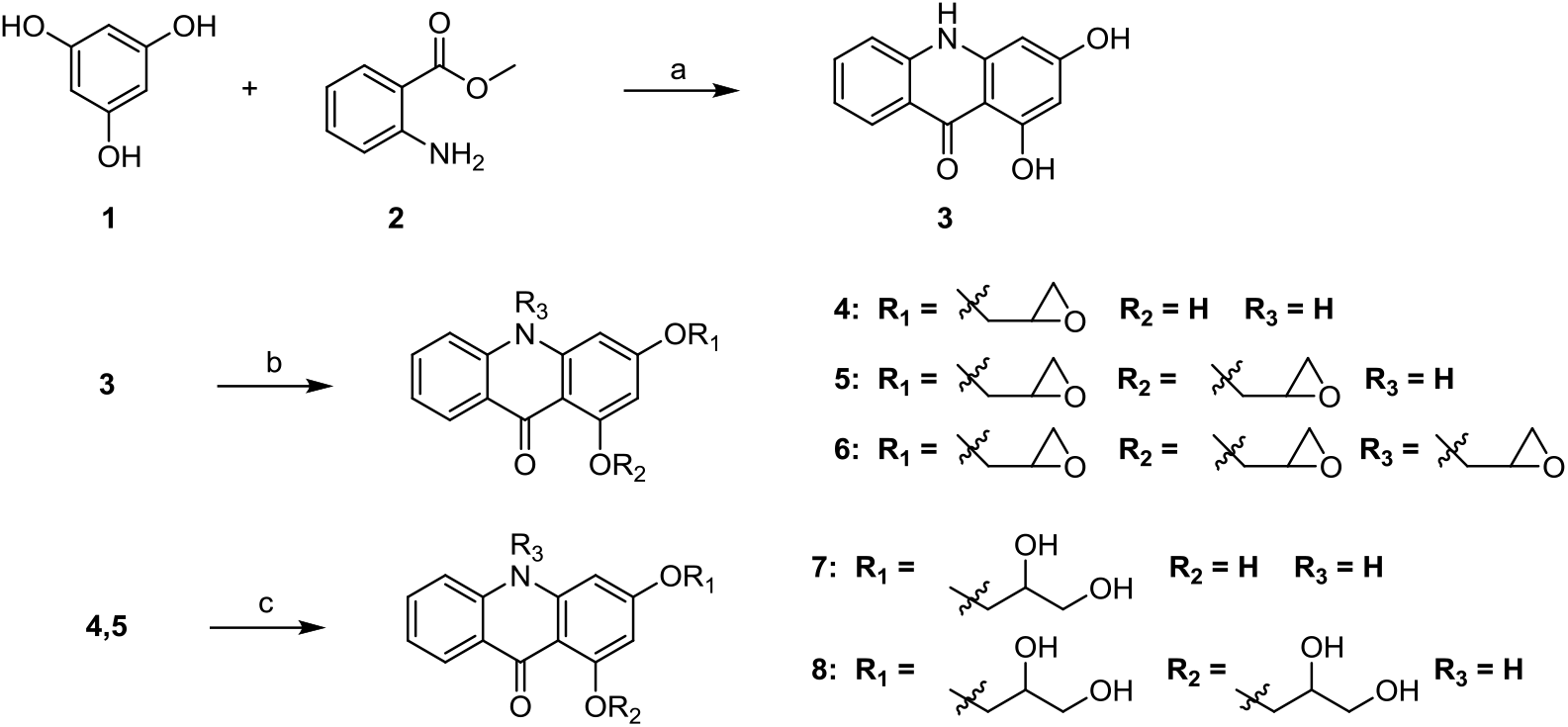
Synthesis of acridone epoxides. Reagents and conditions: (a) PTSA, hexanol, reflux; (b) Epibromohydrin, Cs_2_CO_3_, acetone/DMF, 40-70 °C; (c) 20% NaOH (aq), 70 °C.

In the case of the xanthone-based analogs, phloroglucinol (**1**) was condensed with salicylic acid (**9**) in phosphorous oxychloride with zinc to afford the 1,3-dihydroxy xanthone (**10**) (Scheme 2). Using the same reaction sequence as for the acridones, both the mono and di-epoxy xanthones **11** and **12** were synthesized in one pot in good yield (~80%).

**Scheme 2:**
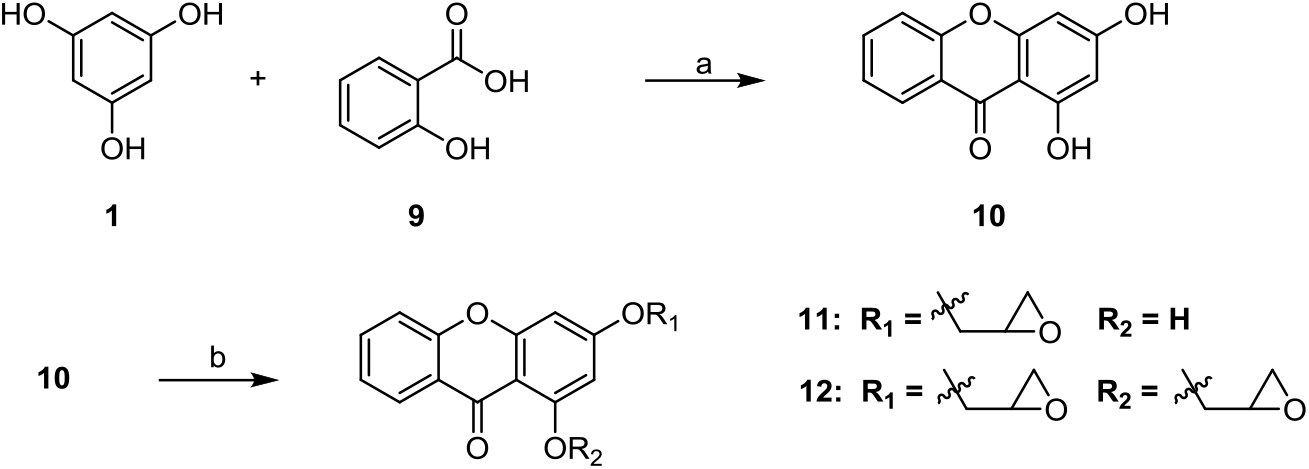
Synthesis of xanthone epoxides. Reagents and conditions: (a) POCl_3_, ZnCl_2_, 70 °C; (b) Epibromohydrin, Cs_2_CO_3_, Acetone/DMF, 40-70 °C.

## Results and Discussion

Anti-bacterial evaluation against both strains of MRSA and MSSA for the acridone and xanthone-based compounds is shown in Table 1. As anticipated, compounds which lack an epoxy group (acridones **3**, **7**, and **8** and xanthone **10**) did not display significant anti-bacterial activity. Similarly, acridone **4** and xanthone **11** with a single epoxy group at the 3 position (R_1_) showed limited antibacterial activity. While bis-epoxy-acridone **5** and the analogous bis-epoxy-xanthone **12** showed good anti-bacterial activity against both strains of MRSA and MSSA, the addition of a third epoxy group yielded no further gains in activity (as shown by compound **6**). Based on these results, it can be concluded that the inclusion of two epoxy groups on either the acridone or xanthone core are critical to obtaining peak antibacterial activity. Vero cell toxicity is also reported in table 1. The two bis-epoxides **5** and **12** show therapeutic indices (TI) that range from 2 to 8 while the mono-epoxide derivative **4** is less discriminant with a TI of 1 to 2. It is interesting to note that highest TI values are associated with activity of tri-epoxide **6** and bis-epoxy xanthone **12** against the MSSA strain, MSA553. This strain, however, is generally more sensitive to antibiotic treatment so the significance of this result is not clear.

**Table 1:** Anti-bacterial Data and Vero Cell Toxicity (MIC: ug/mL).

| Compound | X | R<sub>1</sub> | R<sub>2</sub> | USA400 (MW2) | USA300 (1371) | MSSA (MSA553) | Vero Cell Toxicity |
| --- | --- | --- | --- | --- | --- | --- | --- |
| **3** | NH | OH | OH | >64 | 64 | >64 | NT |
| **4** | NH | Epoxy<sup>a</sup> | OH | 64 | 64 | 32 | 64 |
| **5** | NH | Epoxy | Epoxy | 8 | 16 | 8 | 32 |
| **6** | N-Epoxy | Epoxy | Epoxy | 32 | 16 | 8 | 64 |
| **7** | NH | Diol<sup>b</sup> | OH | >64 | 64 | >64 | NT |
| **8** | NH | Diol | Diol | >64 | >64 | >64 | NT |
| **10** | O | OH | OH | >64 | >64 | >64 | NT |
| **11** | O | Epoxy | OH | >64 | >64 | >64 | NT |
| **12** | O | Epoxy | Epoxy | 8 | 8 | 4 | 32 |

<sup>a</sup>Epoxy denotes 1,2-epoxypropane, <sup>b</sup>Diol denotes 1,2-propanediol, NT = not tested

Given our prior work on xanthones, acridones, and acridines, compounds **5** and **12** were subject to mechanism of action studies.^22–24^ Regulated antisense RNA expression technology was applied to titrate down the expression of a drug target gene and selectively sensitize the bacterial cells to a drug directed to that protein.^20,21^ Both compounds, however, failed to show altered activity to any of the down-regulated essential protein strains in the regulated RNA expression library, including DNA gyrase, DNA topoisomerase, tRNA synthetase, and the fatty acid biosynthesis pathway. This result was somewhat surprising given the history of xanthones and acridones to act as nucleotide binding agents.^25–27^ We cannot rule out the possibility that the bis-expoxides may target other essential proteins involved in cell wall biosynthesis or essential ribosome proteins. Based on the precedence for xanthone-based compounds, such as α-mangostin, to function as nonspecific cytoplasmic membrane disruptors,^14^ the anti-bacterial activity of the bis-epoxides may be due to their ability to covalently bind to bacterial membranes resulting in the loss of cellular integrity. While the specific manner in which the bis-epoxides bind to and disrupt bacterial membranes is beyond the scope of the current investigation, it is interesting to note that the addition of a third epoxide (**6**) does not increase activity. This suggests that the mechanism of action is not simply a function of reactivity, but depends, in part, on specific recognition elements of the target. This concept is further supported by the cell cytotoxicity data that shows the bis-epoxides generally display a better therapeutic index than their mono- and tri-counterparts against the MRSA strains. Although the therapeutic index is modest (~4 fold for both bis-epoxides **5** and **12**), the results indicate bacterial cells may present unique epitopes that could be exploited by further refinement of these reactive heterocyclic scaffolds. Furthermore, a better understanding of their mechanism of action (especially in light of prior work on the nonreactive mangostins) may provide new insight to the design of improved analogs with lower toxicity.

## Materials and Methods

### Chemical synthesis

#### General Experimental Conditions

All chemicals were purchased from Sigma-Aldrich, TCI America, Strem or local vendors and used as supplied unless otherwise noted. All reactions were conducted under an atmosphere of N_2_ or Ar. Thin layer chromatography (TLC) was performed on 0.25 mm hard-layer silica G plates; developed plates were visualized with a hand-held UV lamp or iodine chamber, 10% sulfuric acid or 10% phosphomolybdic acid (PMA) solution. ^1^H and ^13^C NMR spectra were recorded on a Varian 600 MHz or a Varian 400 MHz spectrometer in noted solvent; peaks are reported as: chemical shift (multiplicity, J couplings in Hz, number of protons). High resolution mass spectra (HRMS) were obtained on an Agilent TOF II TOF/MS instrument equipped with either an ESI or APCI interface. Purity was determined based on NMR spectra evaluation and HRMS analysis.

#### 1,3-dihydroxyacridin-9(10H)-one (3)

A mixture of methyl anthranilate (6.6 mmol), phloroglucinol (6.6 mmol), and p-toluene-sulfonic acid (100 mg) in 40 mL of hexanol was heated at reflux for 4 h. The reaction mixture was then cooled, petroleum ether was added, and the mixture stirred well. The product was then filtered and washed with petroleum ether followed by dichloromethane. The resultant solid was recrystallized from ethanol/water to yield 1.28 g (5.63 mmol) of a yellow soild. 85% yield. ^1^H-NMR (DMSO-d_6_): 5.97 (d, *J* = 1.2 Hz, 1H), 6.26 (d, *J* = 0.6 Hz, 1H), 7.22 (t, *J* = 7.5 Hz, 1H), 7.43 (d, *J* = 9.0 Hz, 1H), 7.68 (t, *J* = 7.8, 7.2 Hz, 1H), 8.12 (d, *J* = 8.4 Hz, 1H), 10.49 ( s, 1H), 11.72 ( s, 1H), 14.72 (s, 1H).

#### General procedure for mono and di-epoxy-acridones

1,3-dihydroxyacridin-9(10H)-one (1.1 mmol) and Cs_2_CO_3_ (2.2 mmol) was dissolved in 40 mL of a 1:2 DMF:Acetone mixture. Epibromohydrin (1.23 mmol) was added and the reaction was heated at 40 °C for 16 hours. The reaction was cooled and diluted with water, then extracted with ethyl acetate. The combined organics were washed with brine, dried over Na_2_SO_4_ and concentrated in vacuo. The residue was purified on silica eluting 2:1 Hex:EtOAc to afford both the mono-epoxy and di-epoxy acridone as yellow solids.

### 1-hydroxy-3-(oxiran-2-ylmethoxy)acridin-9(10H)-one (4)

52% yield. ^1^H-NMR (600 MHz, DMSO-d_6_): δ 2.74 (m, 1H), 2.86 (t, *J* = 4.8 Hz, 1H), 3.35 (m, 1H), 3.92 (m, 1H), 4.42-4.45 (dd, *J* = 2.4, 9.0 Hz, 1H), 6.18 (d, *J* = 2.4 Hz, 1H), 6.37 (d, *J* = 2.4 Hz, 1H), 7.26 (t,*J* = 7.7 Hz, 1H), 7.49 (d, *J* = 8.4 Hz, 1H), 7.73 (td, *J* = 8.1, 1.2 Hz, 1H), 8.15 (dd, *J* = 8.1, 1.2 Hz, 1H), 11.9 (s, 1H), 14.22 (s, 1H); HRMS (C_16_H_13_NO_4_) [M-H]^-^: found *m/z* 282.0741, calcd 282.0772

### 1,3-bis(oxiran-2-ylmethoxy)acridin-9(10H)-one (5)

22% yield. ^1^H-NMR (600 MHz, DMSO-d_6_): δ 2.63 (m, 1H), 2.74 (m, 1H), 2.82 (m, 1H), 2.87 (m, 1H), 3.37 (m, 1H), 3.45 (m, 1H), 3.98 (m, 1H), 4.44-4.52 (m, 2H), 4.97 (dd, 1H) 6.33 (d, *J* = 2.4 Hz 1H), 6.67 (t, *J* = 1.8 Hz, 1H), 7.35 (t, *J* = 6.6 Hz, 1H), 7.82 (m, 2H), 8.28 (d, *J* = 6.6 Hz, 1H); HRMS (C_19_H_17_NO_5_) [M+H]^+^: found *m/z* 340.1184, calcd 340.1179.

### 1,3-bis(oxiran-2-ylmethoxy)-10-(oxiran-2-ylmethyl)acridin-9(10H)-one (6)

1,3-dihydroxyacridin-9(10H)-one (2.2 mmol) and Cs_2_CO_3_ (11 mmol) was dissolved in 20 mL of DMF. Epibromohydrin (22 mmol) was added and the reaction was heated at 70 °C for 16 hours. The reaction was cooled and diluted with water, then extracted with ethyl acetate. The combined organics were washed with brine, dried over Na_2_SO_4_ and concentrated in vacuo. The residue was purified on silica eluting 2:1 Hex:EtOAc to yield a yellow soild. 26% yield. ^1^H-NMR (600 MHz, DMSO-d_6_): δ 2.65 (m, 1H), 2.74 (m, 1H), 2.84-2.89 (m, 3H), 3.08 (m, 1H), 3.39 (m, 2H), 3.46 (m, 1H), 4.02 (m, 2H), 4.37 (m, 2H), 4.51 (d, *J* = 11.4 Hz, 1H), 4.88-4.91 (dd, *J* = 4.8, 12.0 Hz, 1H), 6.45 (s, 1H), 6.76 (s, 1H), 7.24 (t, *J* = 7.8 Hz, 1H), 7.68 (m, 2H), 8.20 (d, *J* = 7.2 Hz, 1H); HRMS (C_22_H_21_NO_6_) [M+H]^+^: found *m/z* 396.1543, calcd 396.1520

#### 3-(2,3-dihydroxypropoxy)-1-hydroxyacridin-9(10H)-one (7)

1-hydroxy-3-(oxiran-2-ylmethoxy)acridin-9(10H)-one (0.25 mmol) was dissolved in 15 mL of 20% aqueous NaOH and stirred at 70 °C for 16 hours. The reaction was then cooled and concentrated. The residue was taken up in EtOAc, washed with water, dried over Na_2_SO_4_ and concentrated in vacuo followed by purification on silica eluting 95:5 DCM:MeOH to yield the compound as a yellow solid. 78% yield. ^1^H-NMR (600 MHz, CD_3_OD): δ 3.65-3.82 (m, 2H), 3.9-4.16 (m, 3H), 6.18 (s, 1H), 6.36 (s, 1H), 7.24 (t, *J* = 7.2 Hz, 1H), 7.40 (d, *J* = 8.4 Hz, 1H), 7.67 (t, *J* = 8.4 Hz, 1H), 8.21 (d, *J* = 8.4 Hz, 1H); HRMS (C_16_H_15_NO_5_) [M-H]^-^: found *m/z* 300.0864, calcd 300.0877.

#### 1,3-bis(2,3-dihydroxypropoxy)acridin-9(10H)-one (8)

1,3-bis(oxiran-2-ylmethoxy)acridin-9(10H)-one (0.15 mmol) was dissolved in 15 mL of 20% aqueous NaOH and stirred at 70 °C for 16 hours. The reaction was then cooled and concentrated. The residue was taken up in EtOAc, washed with water, dried over Na_2_SO_4_ and concentrated in vacuo followed by purification on silica eluting 95:5 DCM:MeOH to yield the compound as a yellow solid. 80% yield. ^1^H-NMR (600 MHz, DMSO-d_6_): δ 3.44-3.58 (m, 4H), 3.81 (m, 1H), 4.00 (m, 2H), 4.13 (m, 1H), 4.47 (m, 2H), 6.26 (s, 1H), 6.76 (s, 1H), 7.33 (t, *J* = 7.2 Hz, 1H), 7.79 (t, *J* = 7.2 Hz, 1H), 7.96 (m, 1H), 8.29 (d, *J* = 7.8 Hz, 1H). HRMS (C_19_H_21_NO_7_) [M-H]^-^: found *m/z* 374.1229, calcd 374.1234.

#### 1,3-dihydroxy-9H-xanthen-9-one (10)

Salicylic acid (36.2 mmol), phloroglucinol (55.5 mmol), and zinc chloride (110 mmol) were dissolved in 35 mL of phosphorous oxychloride and stirred at 70 °C for 2 hours. After 2 hours, the mixture was cooled and poured into ice water. The resultant precipitate was collected via filtration, washed with water, and dried. The compound was purified on silica eluting 4:1 Hex:EtOAc to yield the compound as a light yellow solid. 42% yield. ^1^H-NMR (600 MHz, DMSO-d_6_): δ 6.24 (d, *J* = 2.4 Hz, 1H), 6.38 (d, *J* = 2.4 Hz, 1H), 7.45 (t, *J* = 7.2 Hz, 1H), 7.58 (d, *J* = 7.8 Hz, 1H), 7.84 (m, 1H), 8.12 (d, *J* = 8.4 Hz, 1H), 11.09 (s, 1H), 12.80 (s, 1H); HRMS (C_13_H_8_O_4_) [M-H]^-^: found *m/z* 227.0336, calcd 227.0350.

#### General procedure for mono and di-epoxy-xanthones

1,3-dihydroxy-9H-xanthen-9-one (0.88 mmol) and Cs_2_CO_3_ (1.31 mmol) was dissolved in 10 mL of a 1:1 DMF:Acetone mixture. Epibromohydrin (1.23 mmol) was added and the reaction was heated at 60 °C for 16 hours. The reaction was cooled and diluted with water, then extracted with ethyl acetate. The combined organics were washed with brine, dried over Na_2_SO_4_ and concentrated in vacuo. The residue was purified on silica eluting 2:1 Hex:EtOAc to afford both the mono-epoxy and di-epoxy xanthone as pale yellow solids.

#### 1-hydroxy-3-(oxiran-2-ylmethoxy)-9H-xanthen-9-one (11)

40% yield. ^1^H-NMR (600 MHz, DMSO-d_6_): δ 2.71 (m, 1H), 2.85 (t, *J* = 4.2 Hz, 1H), 3.35 (d, *J* = 2.4 Hz, 1H), 3.96 (m, 1H), 4.50 (d, *J* = 11.4 Hz, 1H), 6.42 (s,, 1H), 6.66 (s, 1H), 7.46 (t, *J* = 7.8 Hz, 1H), 7.59 (d, *J* = 8.4 Hz, 1H), 7.86 (t, *J* = 8.4 Hz, 1H), 8.13 (d, *J* = 7.8 Hz, 1H), 12.77 (s, 1H); HRMS (C_16_H_12_O_5_) [M-H]^-^: found *m/z* 283.0632, calcd 283.0612.

#### 1,3-bis(oxiran-2-ylmethoxy)-9H-xanthen-9-one (12)

47% yield. ^1^H-NMR (600 MHz, DMSO-d_6_): δ 2.72 (m, 1H), 2.85 (m, 2H), 3.02 (m, 1H), 3.37 (m, 2H), 3.96 (m, 1H), 4.03 (m, 1H), 4.43 (d, *J* = 11.4 Hz, 1H), 4.50 (d, *J* = 11.4 Hz, 1H) 6.53 (s, 1H), 6.70 (s, 1H), 7.38 (t, *J* = 7.2 Hz, 1H), 7.48 (d, *J* = 8.4 Hz, 1H), 7.73 (t, *J* = 7.2 Hz, 1H), 8.05 (d, *J* = 7.8 Hz, 1H). HRMS (C_19_H_16_O_6_) [M-H]^-^: found *m/z* 341.1052, calcd 341.1020.

### Biological Assays

#### Bacterial strains and growth media

*S. aureus* strains used in this study include MRSA isolates MW2 (USA400) and 1371 (USA300) and MSSA isolate MSA553, and the antisense library of different essential genes.^20,21^ USA300 (1371) and MSA553 isolates were kindly provided by Ds, Richard Goering and Patrick Schlievert, respectively. The *S. aureus* cells were cultured in Trypticase soy broth (TSB) at 37°C with shaking.

#### *S. aureus* susceptibility assay

*S. aureus* strains were grown in TSB at 37°C overnight and were diluted to ~10^5^ CFU/ml as cultures for MIC assays with a 96-well microtiter format. Serial dilutions of the compounds were prepared in Trypticase soy broth with supplement erythromycin (5 μg/ml) (TSB-Erm) in a final assay volume of 100 μl. Fifty μl of 10^5^ CFU/ml bacteria was added to the serially diluted antibiotics. The MIC was considered to be the concentration at which the antibiotic prevented turbidity in the well after incubation for18 h at 37°C.^22^ For antisense RNA expression mutants, the MIC was determined in TSB-Erm in the presence inducer, anhydrotetracycline (100, 250, or 500 ng/ml), to induce antisense RNA expression and downregulate a drug target. The MIC assay was repeated at least three times.

#### Cell culture

Vero monkey kidney epithelial cells (ATCC CCL-81) were cultured in RPMI 1640 medium supplemented with 10% fetal bovine serum(FBS; Invitrogen, CA). Cultures of Vero cells were maintained in a medium containing penicillin (5μg/ml) and streptomycin (100μg/ml) (Invitrogen, CA). Assays were performed in RPMI 1640 medium with different doses of tested compounds.

#### Cytotoxicity assays

The cytotoxicity assay was conducted by measuring LDH release as described.^28^ Briefly, all cells were grown in 96-well plates to 90% confluence. To test the cytotoxicity, monolayer cells were exposed to different doses of tested compounds and incubated for 16 h at 37°C with 5%CO_2_. At the end of the experiment, cell viability was determined by measuring LDH release using the CellTiter 96® Aqueous Non-Radioactive Cell Proliferation Assay (Promega, MI) as per manufacturer’s instructions. Each experiment was repeated three times, and all of the percentage of cell death related to control (no death) were calculated.

## Author Contributions

Padmaja Chittepu completed the chemical synthesis of the antimicrobial drug candidates.

Junshu Yang performed anti-MRSA and anti-MSSA assays and evaluated the mechanism of action of the drug candidates using the RNA expression library technique.

Christine Salomon performed antimicrobial evaluations of the drug candidates.

Adam Benoit prepared samples for biological testing and contributed to writing the manuscript.

Yinduo Ji designed and directed the implementation of the RNA expression library and anti-MRSA and anti-MSSA screening.

David Ferguson conceived and designed the project, wrote the manuscript, and directed the completion of the chemical synthesis and integration of the biological experiments.

## Acknowledgement

This work is supported in part by grant AI078951 (to Y. Ji) from the National Institute of Allergy and Infectious Disease. The M RSA isolate MW2 was obtained through the Network of Antimicrobial Resistance in *Staphylococcus aureus* (NARSA) program supported under NIAID/NIH contract #HHSN272200700055C. We thank John Goodell for helpful comments in the preparation of this manuscript.

